# A Multivariate Comparison of EEG and fMRI to ECoG Using Visual Object Representations in Humans

**DOI:** 10.1101/2022.07.01.498307

**Authors:** Fatemeh Ebrahiminia, Radoslaw Martin Cichy, Seyed-Mahdi Khaligh-Razavi

## Abstract

Today most neurocognitive studies in humans employ the non-invasive neuroimaging techniques fMRI and EEG. However, how the data provided by fMRI and EEG relate exactly to the underlying neural activity remains incompletely understood. Here, we pursued this open question by relating EEG and fMRI data at the level of neural population codes using multivariate pattern analysis. For this, we recorded EEG and fMRI data while participants viewed everyday objects in different viewing conditions, and then related the data to ECoG data recorded for the same stimulus set. The comparison of EEG and ECoG data showed that object category signals emerge swiftly in the visual system and can be detected by both EEG and ECoG at similar temporal delays after stimulus onset. The correlation between EEG and ECoG reduces and onset latencies increase when object representations tolerant to changes in scale and orientation are considered. The comparison of fMRI and ECoG overall revealed a tighter relationship in occipital than in temporal regions, related to differences in fMRI signal-to-noise ratio. Together, our results reveal a complex relationship between fMRI, EEG and ECoG signals at the level of population codes that critically depends on the time point after stimulus onset, the region investigated, and the visual contents used.

## 2. Introduction

Any human cognitive function is realized by complex neural dynamics that evolve both in time and in locus [1, 2] across the brain. No currently available non-invasive neuroimaging technique can resolve such complex brain dynamics both in space and time with high resolution simultaneously. Instead, different techniques are used to pursue one goal or the other. Specifically, fMRI resolves brain activity with high spatial resolution up and below the millimeter scale, but low temporal resolution [3–6]. In contrast, EEG resolves brain activity with high temporal resolution in the millisecond range, but its spatial resolution is limited [7, 8]. However, the data provided by M/EEG and fMRI have a complex and incompletely understood relationship to the underlying neural sources. This complicates their interpretation on their own as well as in combination, introducing methodological uncertainty in our knowledge of the spatiotemporal neural dynamics underlying cognitive functions.

Interpreting the results of fMRI and EEG can benefit from comparison with results from invasive techniques that sample brain activity with high temporal and spatial specificity simultaneously, such as the electrocorticogram (ECoG). A Large body of research has taken this route, relating invasive electrophysiological recordings and EEG or fMRI respectively.

The relationship between EEG signals and invasive electrophysiological recordings has been investigated before mostly in the context of seizure detection [9, 10] and activity localization [11, 12]. Further studies have also explored frequency-band links between the two modalities [9, 13, 14]. An emerging pattern from these studies is that the analytical method used for the comparison [12, 15] and the signal components of EEG and ECoG selected [9, 11] play a key role in achieving a reliable correspondence between the two modalities.

The relationship between fMRI signals and electrophysiological recordings has been investigated in a set of seminal studies in non-human primates during visual processing [16–19]. In the visual cortex, overall BOLD responses reflect LFPs more than the spiking output of the neurons and lower frequency activities [16–19], but the correspondence also varies across brain regions. Studies in humans comparing anatomical and functional correspondence of fMRI and invasive electrophysiological signals show frequency [20–25], region [21, 24, 26], and time specific [27] correspondence, both for task responses and resting state activity [20, 21, 28].

Common to most of the aforementioned studies is that the links between non-invasive neuroimaging signals and ECoG signals were established using univariate analysis. However, the brain codes information in population codes that cannot be captured by univariate analysis. In contrast, multivariate analysis that assesses neural signals at the level of activation patterns can provide complementary information to univariate analysis with potentially higher sensitivity [29–31].

Here, we thus investigated the relationship between EEG, fMRI, and ECoG signals at the level of population codes using multivariate analysis techniques [30, 31]. Our analyses build on previously published ECoG data that was recorded while participants viewed object images from five different object categories, different scales and rotations [32]. We additionally recorded event-related fMRI and EEG responses separately to the same object images. This allowed for a direct content-sensitive comparison between ECoG, EEG, and fMRI signals, detailing how much the correspondence depends on the content (object category) as well as variations in object scale and orientation.

## 3. Materials and Methods

### 3.1. Participants

21 healthy volunteers (13 females, age: mean ± s.d. = 24.61 ± 3.47) with no history of visual or neurological problems participated in this study. All participants went through a health examination before each session and signed a written consent form, declaring that their anonymized data can be used for research purposes. The study was approved by the Iran University of Medical Sciences ethics committee and was conducted in accordance with the Declaration of Helsinki.

### 3.2. Stimulus set

We used a publicly available stimulus set used previously to record ECoG data in humans [32] (for example stimuli see Figure S1 in Liu *et al*. 2009). The stimulus set consisted of 125 grayscale images, with 25 images from each of five object categories: animals, chairs, faces, fruits, and vehicles. Each set of 25 images consisted of 5 different objects under 5 different viewing conditions that were defined by rendered object size and rotation angle: (1) 3° visual angle size, 0° depth rotation, (2) 3° visual angle size, 45° depth rotation, (3) 3° visual angle size, 90° depth rotation, (4) 1.5° visual angle size, 0° depth rotation, and (5) 6° visual angle size, 0° depth rotation.

### 3.3. Data availability

The EEG and fMRI data recorded here are available in Brain Imaging Data Structure (BIDS) [33, 34] format on the OpenNeuro repository [35].

### 3.4. EEG experimental design and stimulus presentation

The duration of the experiment was about one hour. The experiment consisted of 32 runs and participants were allowed to take rest in between every 4 runs. Stimuli were presented on a computer screen subtending 10.69 degrees of visual angle. In each run, all 125 images were presented once in a random order. On each trial, an image was presented for 200 ms followed by 800 ms of blank screen. A fixation cross was presented throughout the experiment, and participants were asked to fixate it. Participants conducted a one-back task on object identity independent of the object’s orientation or size, indicating their response via a keyboard button press. The experiment was programmed using MATLAB 2016 and the Psychophysics Toolbox [36, 37].

### 3.5. EEG acquisition

We recorded EEG signals from 64 sensors with a g.GAMMAsys cap and g.LADYbird electrodes with a g.HIamp amplifier using the 10-10 system. Continuous EEG was digitized at 1,100 Hz without applying any online filters. The electrode placed at the left mastoid was used as the reference electrode. The forehead (FPz) electrode was used as the ground electrode. We further used three electrooculography (EOG) channels to record vertical and horizontal eye movements.

### 3.6. EEG preprocessing

We used EEGLAB [38] for preprocessing. In the first step, we concatenated all EEG data from a given recording and filtered them using a low-pass filter with a cutoff frequency of 40 Hz. We then resampled the signals to 1,000 Hz, re-referenced them to the electrode placed on the left mastoid, and extracted epochs from −100 to +600 ms with respect to the stimulus onset. Last, we used Independent Component Analysis (ICA) to remove components identified to be due to eye blink and movement. For each participant, this procedure yielded 32 preprocessed trials for each of the 125 images.

### 3.7. EEG multivariate pattern classification

We used time-resolved multivariate pattern classification to estimate the amount of information about object category in EEG signals at each time point of the epoch. All analyses were conducted independently for each participant. We classified object categories in a one-versus-all procedure, which is one category (e.g., faces) versus all others (animals, chairs, fruits, and vehicles). We conducted the analyses in three schemes equivalent to the ones used in Liu *et al*. 2009 to evaluate the classification performance for ECoG data [32]: category selectivity, rotation invariance, and scale invariance. The schemes are described in detail below.

The scheme “category selectivity” refers to the training and testing of the classifier during category classification using EEG data for all versions of the stimuli independent of stimulus variations (i.e., images with size of 3° visual angle, and 0°, 45°, and 90° depth rotation, sizes of 1.5° and 6° visual angle, and 0° depth rotation). The analysis was conducted separately for each time point of the epoch and for each category. For each category, there were 800 trials (5 identities × 5 variations × 32 repetitions) resulting in two partitions of 800 vs 3200 trials in the one-vs-other classification scheme. To compensate for the imbalanced class ratio, we randomly selected 800 trials from the 3200 trials belonging to the other four categories. To increase signal-to-noise ratio, we randomly assigned the 800 trials into 150 sets (with five or six trials at each set) and averaged the trials in each set to generate 150 pseudo-trials. We further applied multivariate noise normalization [31]. Then, 149 trials were used to train the linear SVM classifier and data from the left-out trial were used for the testing. We repeated the above procedure 300 times, each time with a different, random selection of 800 trials from the other category class.

The scheme “rotation invariance” refers to classifying object categories across orientation changes. For this, we trained the classifier using EEG data for images with size of 3° visual angle and 0° depth rotation and tested it with EEG data for images of size of 3° visual angle and either 90°or 45° depth rotation, and averaged the results.

Finally, the scheme “scale invariance” refers to classifying object categories across scale changes. For this, we trained the classifier using EEG data for images with size of 3° visual angle and 0° depth rotation and tested it with EEG data for images of size of 1.5° or 6° visual angle and 0° depth rotation, before averaging the results.

For both the rotation and scale invariance schemes, class ratios were imbalanced in the following way: for each category, there were 160 (5 identities × 1 variations × 32 repetitions) trials per participant and thus 160 vs 640 trials in the one-versus-all class sets. To balance the ratio, we randomly selected 160 trials from the set of 640 trials. After applying multivariate noise normalization, we conducted leave-one-trial-out classification. We repeated the procedure 150 times and averaged the results.

### 3.8. ECoG experimental design and procedure

A detailed description of the ECoG data recording is available in the original publication [32]. Here, we provide a short summary. The ECoG data were recorded from 912 subdural electrodes implanted in 11 participants (6 male, 9 right-handed, age range 12–34 years) for evaluation of surgical approaches to alleviate a resilient form of epilepsy. Participants were presented with images of the same set as described above in pseudorandom order. On each trial, an image was presented for 200 ms, followed by 600 ms of blank screen. Out of the 912 electrodes evaluated, 111 (12%) showed visual selectivity. These electrodes were none uniformly distributed across the different lobes (occipital: 35%; temporal: 14%; frontal: 7%; parietal: 4%). The areas with the highest proportions of selective electrodes were inferior occipital gyrus (86%), fusiform gyrus (38%), parahippocampal portion of the medial temporal lobe (26%), midoccipital gyrus (22%), lingual gyrus in the medial temporal lobe (21%), inferior temporal cortex (18%), and temporal pole (14%) (30).

### 3.9. ECoG multivariate pattern classification

Liu *et al*. 2009 [32] used an SVM classifier with a linear kernel to calculate category selectivity, rotation, and scale invariance. They built a neural ensemble vector that contained the range of the signal (max(x)-min(x)) in individual bins of 25 ms duration using 11 selective electrodes and the 50–300 ms interval post stimulus onset. They tested for significance through randomly shuffling the category labels. Here, rather than re-analyzing the data, we reproduced the results as reported in Liu *et al*. 2009 [32].

### 3.10. FMRI experimental design and stimulus presentation

Each participant completed one session consisting of 8 functional runs of data recording. During each run, the same set of images used in EEG and ECoG were presented once in a random order. On each trial, the image was presented for 200 ms followed by a 2,800 ms blank screen. In each run, there were also 30 null trials during which a gray screen with a fixation cross at the center was presented for 3,000 ms. As in the EEG design, participants conducted a one-back task on object identity independent of the object’s orientation or size. The stimulus set was presented to the participants through a mirror placed in the head coil.

### 3.11. FMRI acquisition

We recorded fMRI data using a Siemens Magnetom Prisma 3 Tesla with a 64-channel head coil. MRI head cushions and pillows were used to comfort participants and minimize head movements. To dampen the scanner noise, all participants were provided with a set of earplugs. Structural T1-weighted images were acquired at the beginning of the session for all participants (voxel size = 0.8×0.8×0.8 mm, slices per slab = 208, TR = 1,800 ms, TE = 2.41 ms, flip angle = 8°). We obtained functional images using an EPI sequence with the whole brain coverage (voxel size =3.5×3.5×3.5 mm, TR = 2,000 ms, TE = 30 ms, flip angle = 90°, FOV read = 250 mm, FOV phase = 100%).

### 3.12. FMRI preprocessing and regions of interest extraction

SPM8 [39] and MATLAB 2016 were used to preprocess functional and structural magnetic resonance images. The slices of the functional images were temporally realigned to the middle slice. Then, the volumes of the whole session were aligned to the first volume and co-registered to the structural images.

We used a General Linear Model (GLM) to estimate condition-specific activations. For each run, the onsets of image presentation entered the GLM separately as regressors of interest and were convolved with a canonical hemodynamic response function. We included motion regressors as regressors of no interest. This procedure yielded 125 parameter estimates per run, indicating the responses of each voxel to the 25 different objects presented.

We used Freesurfer package [40] with Destrieux and Desikan-Killiany atlases to extract six regions of interest (ROIs) from surface-based parcellations derived from the T1 images. We used 6 visual ROIs: occipital-inferior, fusiform, lingual, parahippocampal, inferior temporal, and pole temporal cortex. We constructed a binary mask for each region based on the individuals’ structural image parcellation. We then selected voxels in the fMRI data overlapping with the binary mask for further ROI-specific analysis.

### 3.13. FMRI multivariate pattern classification

We used multivariate pattern classification to estimate category selectivity, rotation invariance, and scale invariance at each participant and ROI. Equivalent to the EEG and ECoG analysis, we conducted three analysis schemes: category selectivity, scale invariance, and rotation invariance. We thus do not describe the general setup again, but restrict ourselves to particulars to the fMRI analysis.

In the “category selectivity” scheme, we employed a leave-one-run-out cross-validation procedure to train and test the classifier. There were 8 repetitive functional runs of data recording. For each category, there were 175 trials (5 identities × 5 variations × 7 repetitions) in the training set, resulting in two partitions of 175 vs. 700 trials in the one-vs-other classification scheme. To compensate for the imbalanced class ratio, we randomly selected 175 trials from the 700 trials belonging to the other four categories. We tested the classifier with the left-out run to discriminate between each category and all other four categories. We repeated the random selection of trials 300 times. The classification performances reported here are the average values across the repetitions.

In scale and rotation invariance analyses, for each category and participant, there were 40 (5 identities × 1 variations × 8 repetitions) trials and thus 40 vs. 160 trials in the one-versus-other class sets. To balance the number of trials for two classes (one category vs. the other four categories) in the training set, we randomly selected 40 trials from the other four categories (160 trials). We tested the classifier using a one-trial-out procedure, repeated these steps 300 times and averaged the results.

### 3.14. Relating EEG, fMRI, and ECoG data

We used two principled ways to relate EEG, fMRI and ECoG data to each other: in terms of classification performance resulting from multivariate pattern analysis, and in terms of similarity relations between multivariate activation patterns using representational similarity analysis. We detail each approach below.

#### 3.14.1. Comparison using classification performance

Relating EEG and ECoG, in general we compared their signals by comparing their classification time courses over time with each other. To enable this step, for EEG, we averaged the EEG classification time courses across all participants, yielding one classification time course for each analysis scheme. For ECoG, we used the classification time courses as provided by Liu *et al*. 2009 [32].

In detail, we compared EEG and ECoG signals for each classification scheme in two ways. In the first analysis, we calculated the correlation (Spearman’s R) between the complete accuracy time courses. The second analysis was more fine grained in that it was time-resolved and depended on category-specific time courses. For every time point, we aggregated the classification accuracies for each of the five categories into a 1 (time point) *5 (category) vector. We did so independently for ECoG and EEG, and then compared the vectors using correlation (Spearman’s R). This yielded one correlation value at every millisecond.

Relating fMRI and ECoG, we considered each classification scheme and ROI separately. To enable this step, for fMRI, we averaged the classification results across participants. For ECoG, we again used the classification time courses as provided by Liu *et al*. 2009 [32].

#### 3.14.2. Comparison using Representational Similarity Analysis

We used representational similarity analysis (RSA) [41–43] to relate fMRI, EEG, and ECoG to each other. RSA abstracts from the incommensurate signal spaces of the neuroimaging modalities into a similarity space defined through representational similarity of activation patterns between experimental conditions in each modality. As the 125 experimental conditions were the same in fMRI, EEG, and ECoG, their similarity space is directly comparable. RSA uses dissimilarity rather than similarity by convention which otherwise leaves the rationale of the procedure untouched.

Dissimilarities between conditions are stored in representational dissimilarity matrices (RDMs). RDMs are indexed in rows and columns by the experimental conditions compared, are symmetric across the diagonal, and the diagonal is undefined. Here defined by our stimulus set, the RDMs are of dimensions 125*125.

We constructed RDMs using 1-Spearman’s R as a dissimilarity measure in each case. In detail, for EEG, for every millisecond of the epoch separately, we constructed one RDMs from sensor-level activation patterns. For ECoG, we calculated two different RDM versions. The first version we call “ECoG RDMs”: for every time point, we used data aggregated across all participants in all electrodes (Figure S1) to calculate pairwise dissimilarities between experimental conditions. The second we call “ECoG regional RDMs”. Here, we differentiated between electrodes by ROIs, calculating RDMs separately for each ROI rather than across all electrodes. For fMRI and for each participant and ROI, we constructed RDMs based on ROI-specific beta activation patterns.

### 3.15. Assessing how fMRI signal-to-noise ratio affects the correspondence between fMRI and ECoG

In fMRI recordings, the signal to noise ratio (SNR) is known to differ across the brain. For example, due to artifacts related to the closeness of the ear canal, temporal brain regions often suffer from SNR loss [26, 44–46]. We therefore explored if there is a significant relation between the SNR in a given ROI and the relationship between ECoG and fMRI in the classification performance based analysis. To this end, for fMRI, we averaged classification accuracies across participants at each region. Then, we calculated Spearman’s correlation between mean classification accuracies for ECoG and MRI. This gave one correlation coefficient per region (i.e., a 1×6 vector). Finally, the relation between ECoG-fMRI correlation and SNR was obtained using Spearman’s correlation. We used the upper bound of the noise-ceiling as defined in [47] as a proxy for SNR in the fMRI data.

### 3.16. Inferential analysis

We used non-parametric statistics that do not make assumptions about the distribution of the data for inferential analysis.

#### Peak latency

The time for peak decoding accuracy was defined as the time where the decoding accuracy was the maximum value.

We used bootstrap resampling of participants (21 participants with 10,000 repetitions) to examine whether peak latencies between EEG and ECoG classification time courses were significantly different.

#### Onset latency

We defined onset latency as the earliest time where performance became significantly above chance for at least 15 consecutive time-points. We used the non-parametric Wilcoxon signrank test (one-sided) across participants to determine time-points with significantly above chance decoding accuracy. To adjust p-values for multiple comparisons (i.e. across time), we further applied the false discovery rate (FDR) correction.

We further used bootstrap resampling of participants (21 participants with 10,000 repetitions) to examine whether onset latencies between EEG and ECoG classification time courses were significantly different.

To assess the statistical significance of RDM-to-RDM Spearman’s R correlation, we used a random permutation test based on 10,000 randomizations of the condition labels; where required, the results were FDR-corrected at p < 0.05.

#### Overall Correlation coefficients across time

We used bootstrap resampling of participants (21 participants with 10,000 repetitions) to investigate whether EEG-ECoG RDM correlations were significant.

#### Classification performance

We used bootstrap resampling of participants (21 participants with 10,000 repetitions) to examine if decoding accuracies in fMRI were significantly higher than chance level.

## 4. Results

### 4.1. Comparing time courses of EEG and ECoG-based object category classification

We determined similarities and differences with which EEG and ECoG reveal the temporal dynamics of visual category representations in the human brain. For this, we investigated time-resolved object category sensitivity in neural signals and their robustness to changes in viewing conditions using multivariate pattern analysis. In particular, we classified object category in three different classification schemes (Figure 1A-C). In the first analysis, we classified the object category based on neural signals lumped together for objects of all sized and rotations, indexing overall object category selectivity of the neural signals (analysis from here on termed “category selectivity”). In the second and third analyses, we classified the object category across changes in rotation and scale of the objects. This indexes category selectivity of the neural signals tolerant to changes in viewing conditions.

**Figure 1.**
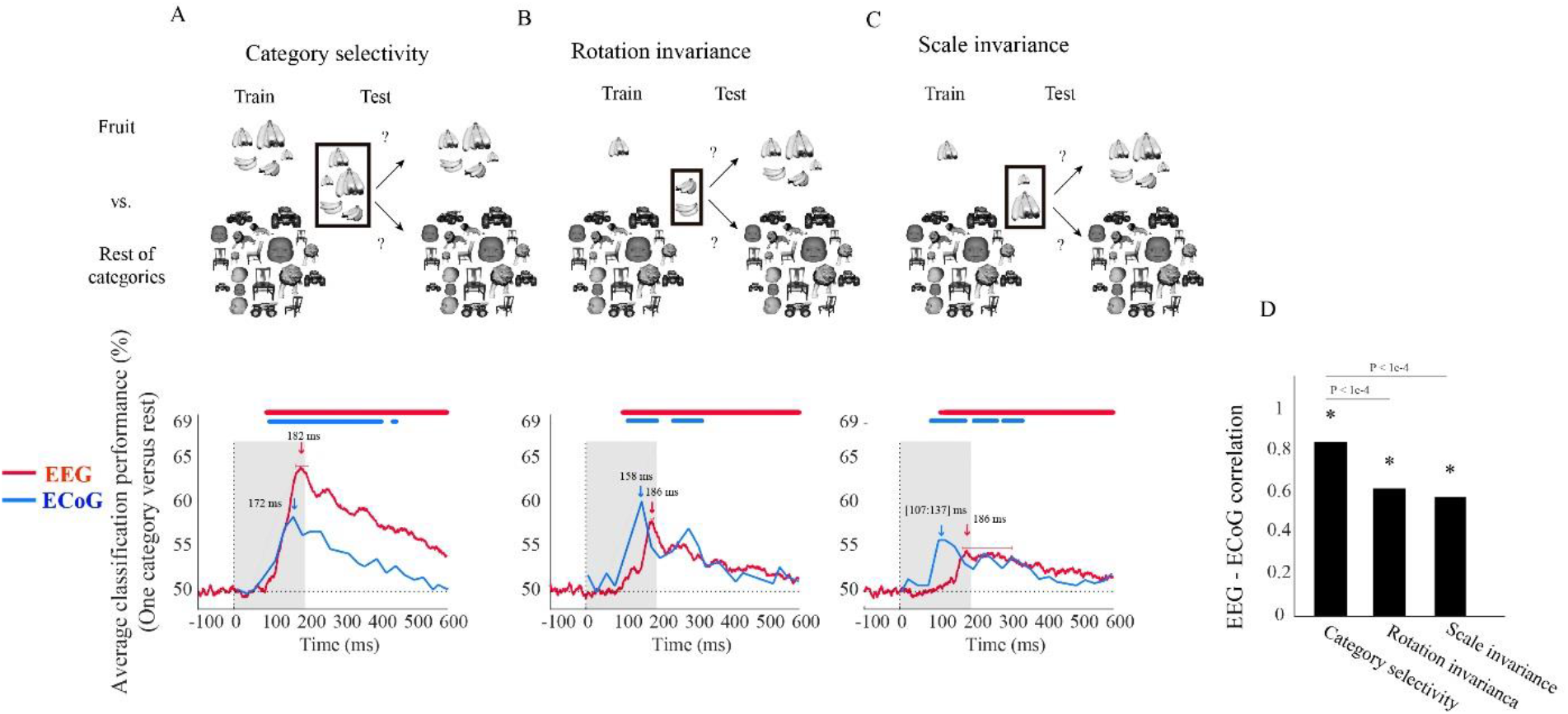
EEG and ECoG average classification performance (one category versus rest) assessing. **(A)** category selectivity, **(B)** rotation invariance, and **(C)** scale invariance over time. The sketches in the top row visualize the rationale of the three classification schemes (i.e., determining category selectivity, rotation invariance, and scale invariance) by showing example images for data entering the classifications. The bottom row shows the classification results. The red curves show EEG classification performance averaged over 5 categories and participants (N = 21). The blue curves show the mean ECoG classification performance over 5 categories using 11 selective electrodes and IFP power in bins of 25 ms. The horizontal lines above the curves indicate significant time points (EEG: bootstrap test of 21 participants with 10,000 repetitions, FDR-corrected at p < 0.05; ECoG: as in the Liu *et al*. 2009, i.e., permutation test over stimuli, not corrected, for details on peak latency comparison see table S1). (**D)** Correlation between EEG and ECoG classification time courses. Stars show significance, and the values above the bar charts indicate the p-values of the differences between the correlation coefficients (bootstrap test of 21 participants with 10,000 repetitions, for more details see table S2).

We begin the comparison of EEG vs ECoG signals at a coarse level by inspecting the shape of the grand average classification results from the above-mentioned classification schemes. We found that object category was robustly classified from neural signals in both EEG and ECoG in all three analysis schemes (Figure 1A-C). In each case, we observed a rapid rise of classification accuracy after image presentation to a peak followed by a gradual decline, with no significant differences in the onset latencies for EEG and ECoG (bootstrap test of 16 participants with 10,000 repetitions and signrank test, all p > 2e-1). This shows that both measurement techniques equally reveal the swift emergence of object category signals in the visual system.

Based on this finding, we quantitatively compared the dynamics with which object category signals become visible in EEG and ECoG for similarity. For this, we correlated their respective time courses across time for each analysis scheme separately. There was a significant relationship in all three cases (Figure 1D), being highest in the general category selectivity analysis and reduced for the scale-and rotation tolerant analysis (bootstrap test of 21 participants with 10,000 repetitions; for detail see table S2).

Besides similarities, we also observed two marked differences between the classification curves for EEG and ECoG. For one, the peak latency for ECoG was significantly shorter than that of EEG in all three analysis schemes (bootstrap test of 21 participants with 10,000 repetitions, for detail see table S1). Second, peak classification accuracy was significantly higher for EEG than for ECoG in the category selectivity analysis (Figure 1A; bootstrap test of 21 participants with 10,000 repetitions, p < 9.9e-4), but lower in the rotation and scale invariant analysis (Figure 1B and C; difference not significant, p > 9.3e-1). This refines the point that neural signals recorded with ECoG and EEG differ in that the ECoG signals reflect visual object representations tolerant to changes in visual processing proportionally more strongly than the EEG signals.

We continue the comparison of EEG vs ECoG signals at a finer level by inspecting the shape of the classification results at each object category separately (Figure 2) to refine the conceptual resolution of the analysis. Consistent with the previous analyses, we observed that each single category was robustly classified from both EEG and ECoG data in all three analysis schemes (Figure 2B). Further consistent with the previous analysis, we found significantly shorter peak latencies for ECOG than for EEG signals for all categories except fruits in the category selectivity scheme, but the difference was not significant for category selectivity under rotation and scale changes (bootstrap test of 21 participants with 10,000 repetitions; for details see Table S3).

**Figure 2.**
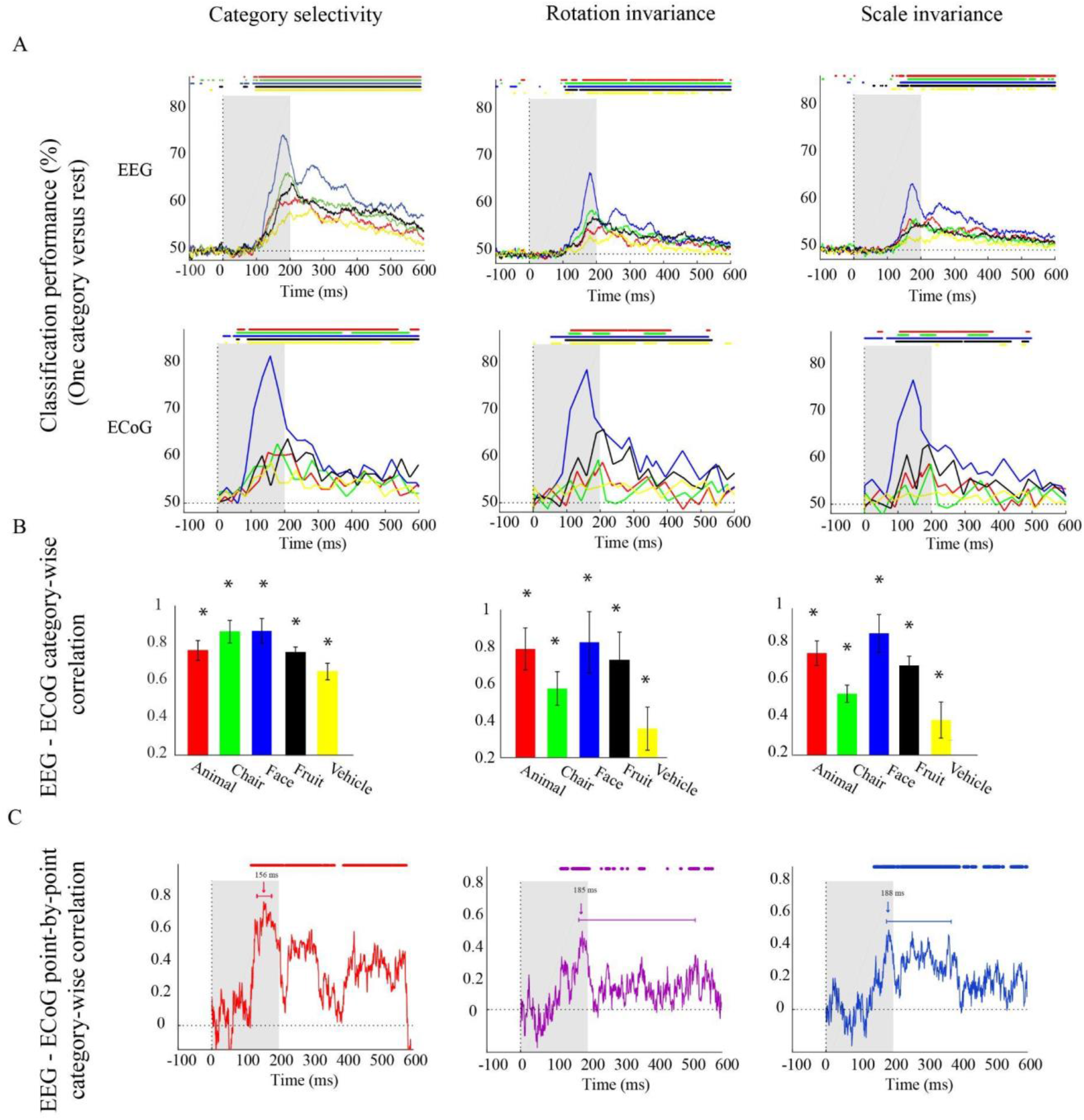
EEG and ECoG category-specific classification performances over time. **(A)** Classification performances across all channels of EEG averaged over 21 participants, and ECoG classification performances using 11 selective electrodes and IFP power in bins of 25 ms. The horizontal lines above the curves indicate significant time points (EEG: bootstrap test of 21 participants with 10,000 repetitions, FDR-corrected at p < 0.05, ECoG: permutation test over stimuli, not corrected). **(B)** EEG - ECoG category-wise correlation. Bar charts show the correlation coefficients between the classification time courses performances of EEG and ECoG for each category separately for the two modalities. The bars are color-coded, and the stars indicate significance (bootstrap test of 21 participants with 10,000 repetitions). **(C)** EEG - ECoG time-resolved point-by-point category-wise correlation. For each time point, we compared classification performance across categories between EEG and ECoG. At each time point, accuracies for categories were compared between EEG and ECoG. The horizontal lines above the curves indicate significant time points (bootstrap test of 21 participants with 10,000 repetitions, FDR-corrected at p < 0.05). For ECoG the results are presented as reported in Liu et al. 2009. We did not analyze the data; we reproduced the figures in Liu *et al*. 2009 [32].

With respect to onset latencies, ECoG had a significantly earlier onset compared to EEG for all object categories under the category selectivity scheme; but this was not the case under scale and orientation variations (bootstrap test of 21 participants with 10,000 repetitions; p-values are given in Table S4). However, in the grand average analysis (Figure 1A-C) there were no significant differences between ECoG and EEG onset latencies in none of three schemes.

The finer level of inspection at the level of single categories allowed us also to compare time courses of classification between different categories. We found that in both EEG and ECoG and across all the three classification schemes, the face category was discriminated earlier and had a significantly higher peak amplitude compared to other categories (Figure 2A; bootstrap test of 21 participants with 10,000 repetitions, all Ps < 1e-3). This shows that the consistency between ECoG and EEG also depends on the content, i.e. the category used to probe the visual system.

To further quantitatively determine similarities between EEG and ECoG at this finer level, we conducted two analyses. First, analogous to the analysis above, we correlated the respective time courses for each category and analysis scheme separately (Figure 2B). We found a positive relationship in all cases (bootstrap test of 21 participants with 10,000 repetitions; for details see Table S5). Our results show that after introducing changes in size and orientation, the correlation between EEG and ECoG decoding accuracies was significantly reduced for the chair and the vehicle category (Figure 2B; bootstrap test of 21 participants with 10,000 repetitions, all Ps < 4e-3). Second, we related EEG and ECoG by correlating the vector of category-specific classification results for each time point separately. Again, we found positive relationships in all cases. Highest similarity and shortest onset latencies were observed in the general category selectivity analysis compared to the scale and rotation tolerant analyses (bootstrap test of 21 participants with 10,000 repetitions; for detail see Tables S6, S7). Consistent with the grand average analysis (Figure 1D), we see that the similarity between EEG and ECoG is reduced and delayed after introducing variations in object scale and orientation.

### 4.2. Comparing EEG and ECoG by representational similarity

In the previous sections, we compared EEG to ECoG in terms of object category classification time courses in classification schemes that assess the effect of changes across scale and invariance.

Here, we use representation similarity analysis (RSA) [41–43] to compare EEG and ECoG representations across time and selected brain regions, regardless of the object category and changes in scale and orientation.

We conducted two analyses defined by the way ECoG data were aggregated in representational dissimilarity matrices (RDMs). In the first analysis (Figure 3A), ECoG RDMs were calculated from data of all ECoG electrodes (called here “ECoG RDMs”), giving a spatially unspecific large-scale statistical summary of representational relations in the ECoG signal. We observed two time periods at which EEG and ECoG representations were significantly similar (i.e., 154 to 222 ms, and 274 to 376 ms) (Figure 3A). We note that the first time period overlaps with both EEG and ECoG peak decoding accuracies at 182 ms and 172 ms, respectively (Figure 1A).

**Figure 3.**
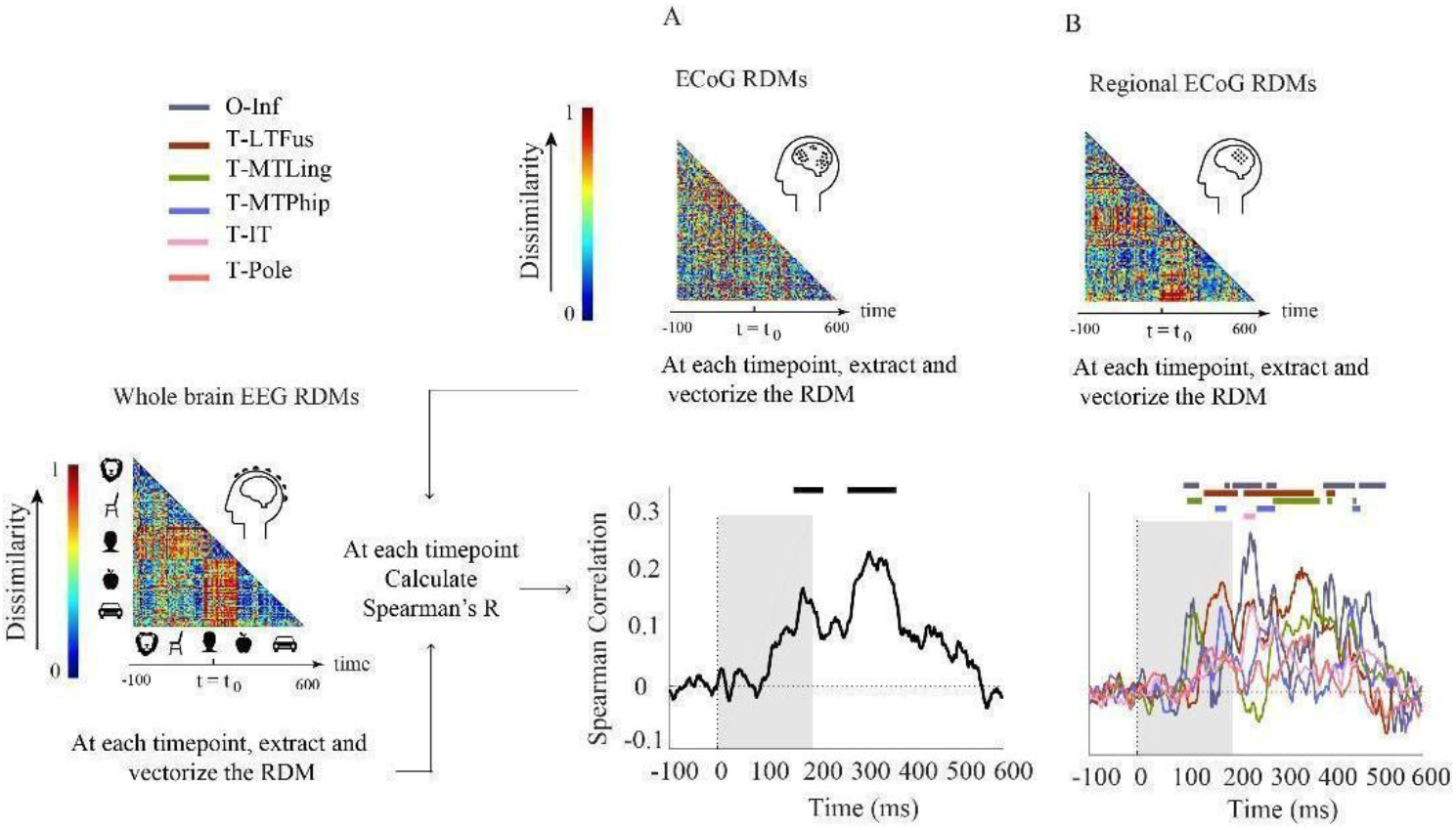
Comparison of EEG and ECoG by representational similarity. We compared the EEG RDM (mean over participants) to **(A)** the ECoG RDM, constructed using all electrodes from all participants and **(B)** the regional ECoG RDM, constructed from electrodes limited to specific ROIs. The horizontal lines above the curves indicate significant time points (permutation test over stimuli, 10,000 iterations, FDR-corrected at p < 0.05). The ROIs include occipital-inferior (O-Inf), fusiform (T-LFus), lingual (T-MLing), parahippocampal (T-MTPhip), inferior-temporal (T-IT), and pole-temporal cortex (T-Pole).

In the second analysis (Figure 3B), we limited the analysis to electrodes in predefined visual ROIs, yielding ECoG RDMs that enable spatially specific assessment of the ECoG data per region (called here “Regional ECoG RDMs”).

We observed a significant representational similarity between EEG and ECoG in occipital-inferior, lingual, fusiform, parahippocampal, and inferior-temporal regions (Figure 3B). Comparison of the results in Figure 3A and 3B suggests how each region may have contributed differently to the overall EEG-ECoG correlation across time. Occipital-inferior and lingual regions are the regions that show the first significant correlation value at 100 and 106 ms respectively. Then the fusiform region shows first significant correlations at 119 ms, followed by the parahippocampal and temporal-inferior regions at 177 and 230 ms. Furthermore, we find that there were significant differences between the regions’ latencies of the first peak; occipital-inferior and lingual peaked at 118 and 116 ms, followed by the fusiform (178 ms), parahippocampal (142 ms) and temporal-inferior region (186 ms) (bootstrap test of 21 participants with 10,000 repetitions, for details see Table S8). Together, this suggests that the fusiform region contributes to the overall pattern of results more than the other regions investigated here (Figure 3A vs. red curve in Figure 3B).

### 4.3. Comparing fMRI and ECoG through object category classification performance

We began the comparison of fMRI with ECoG signals by evaluating the results of category classification in the same three classification schemes as used previously. We focused the analysis on the same six ROIs as in the ECoG-EEG comparison: occipital-inferior, fusiform, lingual, parahippocampal, inferior-temporal, and pole-temporal cortex. Focusing on spatial rather than temporal specificity, for ECoG, we used the classification performance in the time range of 50 to 300 ms after stimulus onset as calculated in Liu *et al*. 2009 [32], as this period captures the visual systems first response to external stimuli as detected by electrophysiological modalities.

The results (Figure 4) revealed a nuanced picture showing both similarities and differences between fMRI and ECoG, depending on category classified, region, and analysis scheme.

**Figure 4.**
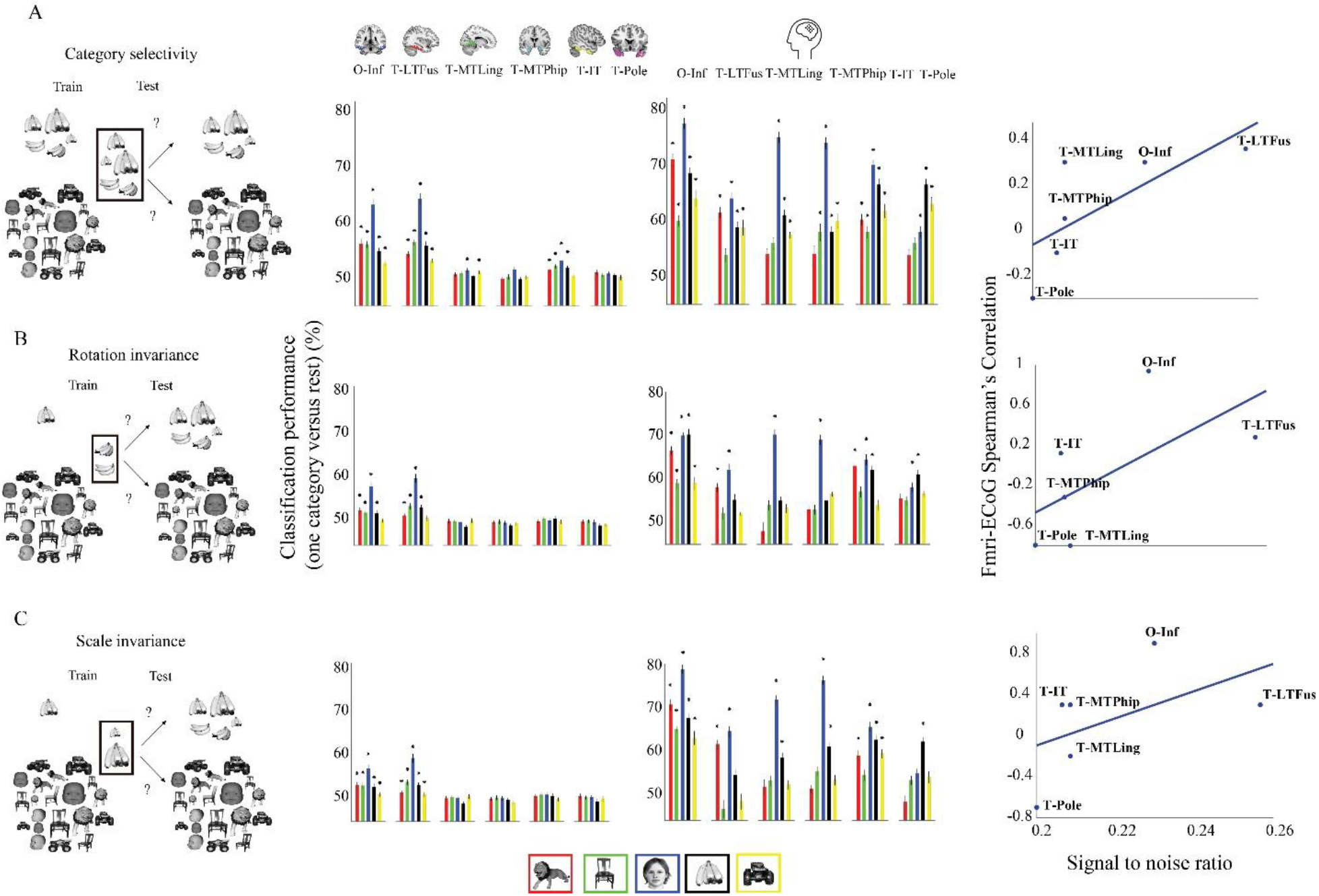
Classification performance (one category vs rest) for the classification schemes. **(A)** category selectivity, **(B)** rotation invariance, **(C)** and scale invariance. First column: cartoon visualization of stimuli for data used to train and test the classifier in each analysis scheme. Second column: region-specific classification results for (average performances across 21 participants). Third Column: region-specific classification results for ECoG. Error bars indicate one standard deviation, stars above bars indicate statistical significance (fMRI: bootstrap test of 21 participants with 10,000 repetitions, FDR-corrected at p < 0.05. ECoG: permutation test over stimuli, not corrected). Fourth Column: Correlation between signal-to-noise ratio in fMRI and ECOG classification results in the category selectivity scheme across regions. The ROIs include occipital-inferior (O-Inf), fusiform (T-LFus), lingual (T-MLing), parahippocampal (T-MTPhip), inferior-temporal (T-IT), and pole-temporal cortex (T-Pole).

First, decoding accuracies in ECoG were significantly higher than that of fMRI for all categories and for all three classifications schemes (bootstrap test of 21 participants with 10,000 repetitions, FDR-corrected at p < 0.05). Further, compared to ECoG, fewer categories could be decoded in fMRI in the orientation and scale invariance scheme, suggesting that some of the information relevant for invariant object recognition was captured less well by the fMRI signal in our experiment. Second, in all three analysis schemes, animal, chair, face, fruit, and vehicle categories were decoded significantly in both fMRI and ECoG in the occipital-inferior region. Third, in the fusiform region, when examining results for the rotation and scale invariance analysis schemes, we observed that all categories could be decoded using fMRI, but in ECoG the chair category could not be decoded.

This complicated picture across regions led us to ask about underlying factors that could explain those results. We hypothesized that the differential signal-to-noise-ratio (SNR) of fMRI across regions might play a role. To investigate, we plotted SNR in fMRI against the similarity in result patterns for fMRI and ECoG for each region and classification scheme (Figure 4, last column). Correlating SNR with fMRI-ECoG similarity values we observed strong positive relationships (bootstrap test of 21 participants with 10,000 repetitions, all Ps < 0.01): regions with low SNR in fMRI such as the ventral occipital temporal and inferior lateral temporal region showed low correlation in classification results patterns with ECOG, whereas regions with high SNR in fMRI such as the occipital inferior and temporal fusiform region showed high correlations. This shows that SNR differences in fMRI across regions strongly contribute to whether positive relationships with ECoG can be established.

### 4.4. Comparing fMRI to ECoG by representational similarity

We further studied the relationship between fMRI and ECoG in a region-specific way using RSA (Figure 5) in a similar way as delineated above for the relationship between EEG and ECoG (Figure 3). In detail, we compared regional fMRI RDMs with regional ECoG RDMs across time.

**Figure 5.**
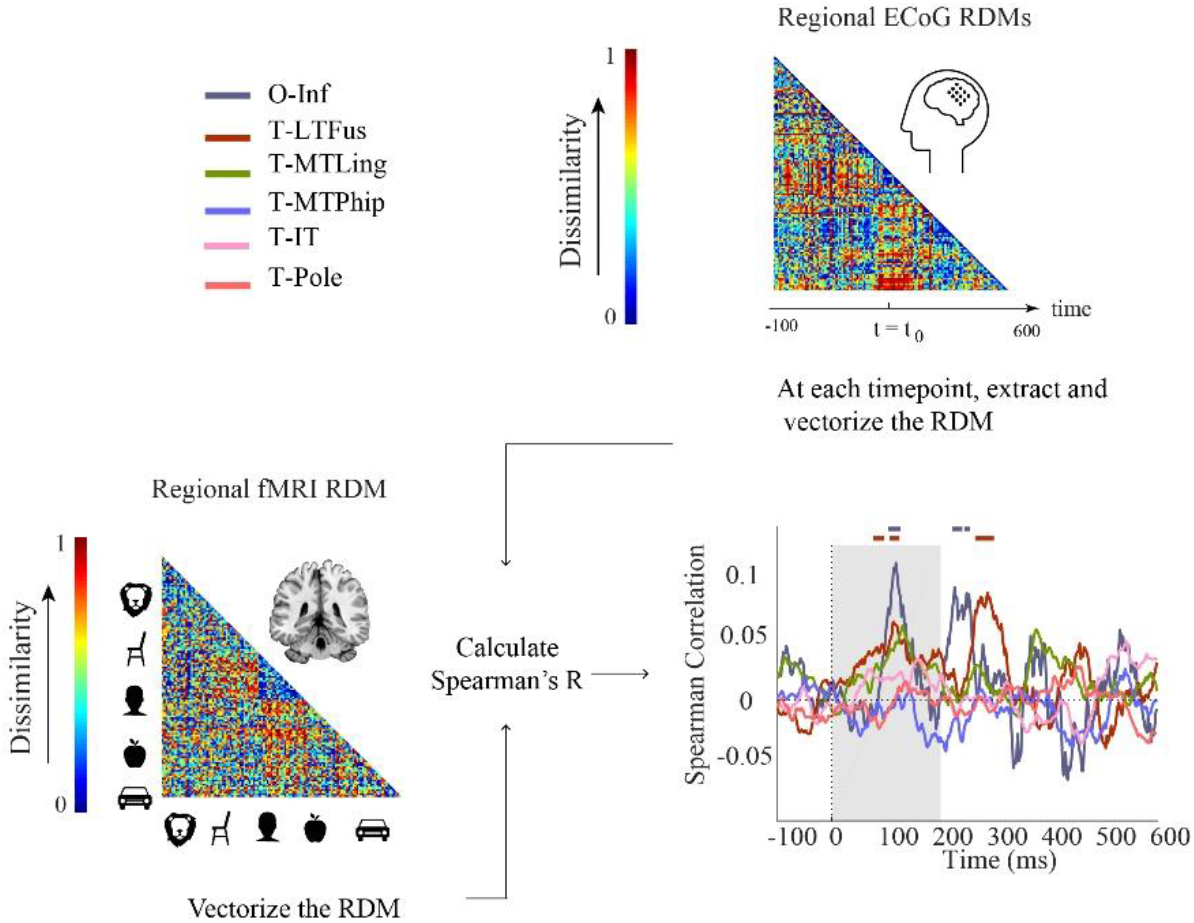
Comparison of fMRI and ECoG in representational similarity by cortical regions. For each participant, fMRI RDMs were calculated using 1-Spearman’s correlation. We then compared the mean fMRI RDM to the ECoG regional RDMs. The horizontal lines above the curves indicate significant time points (permutation test over stimuli, 10,000 repetitions, FDR-corrected at p < 0.05). The ROIs include occipital-inferior (O-Inf), fusiform (T -LFus), lingual (T-MLing), parahippocampal (T-MTPhip), inferior-temporal (T-IT), and pole-temporal cortex (T-Pole).

We found representations as measured by fMRI and ECoG to be similar in occipital-inferior and fusiform regions (Figure 5). The correlation time courses indicating representational similarity showed two peaks around 100 and 250 ms for the occipital-inferior region and around 100 and 300 ms for the fusiform region (for details see Table S9). This suggests that two temporally distinct neural processes occur in those regions, possibly related to feed-forward and feedback processing.

### 4.5. Comparing fMRI to EEG by representational similarity

Finally, aiming to reveal neural dynamics in humans resolved both in space and time using non-invasive techniques only, we used fMRI-M/EEG representational fusion [48–50]. In detail, we compared regional fMRI RDMs with EEG RDMs across time (Figure 6).

**Figure 6.**
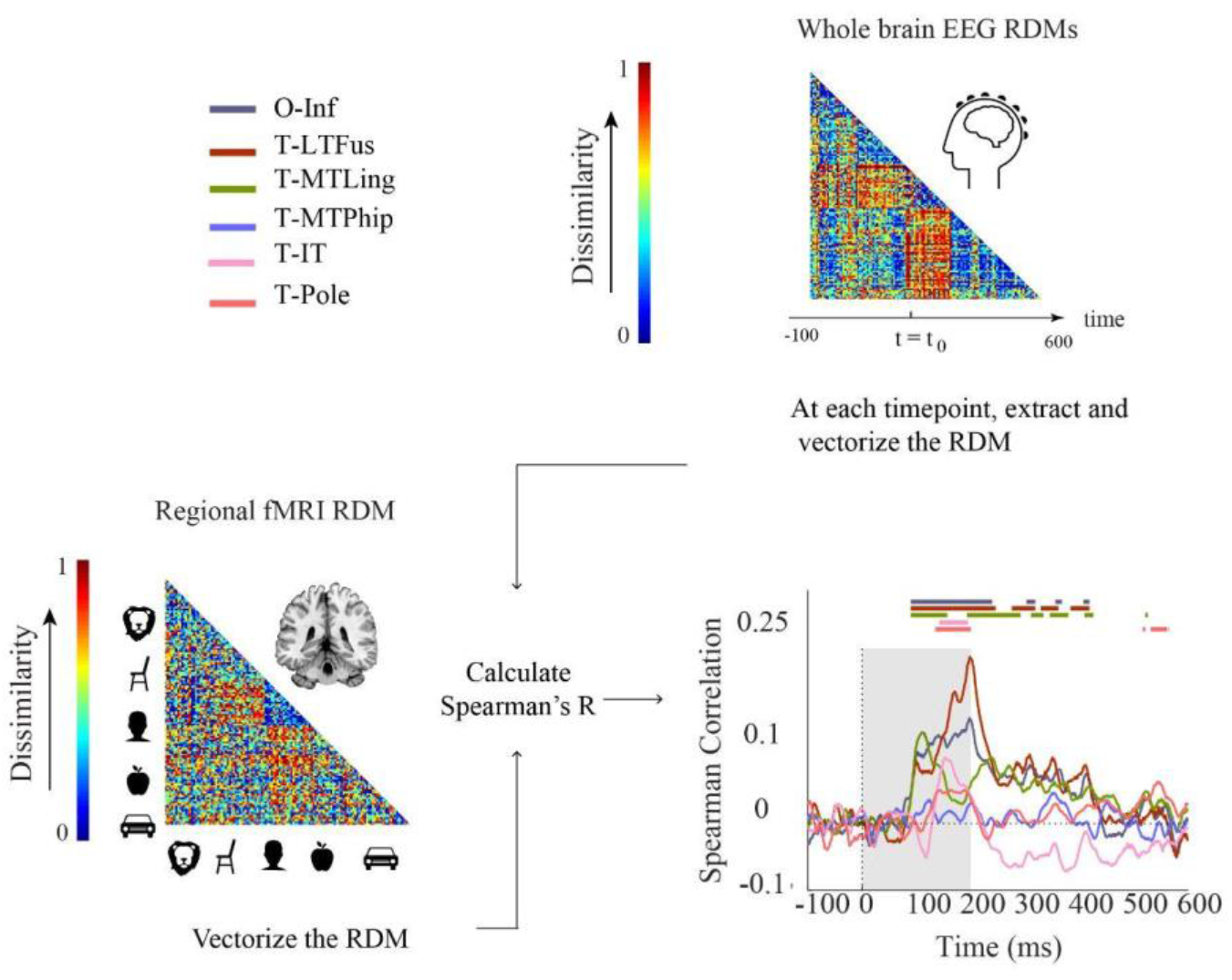
Comparing fMRI and EEG by representational similarity. For each participant, EEG and fMRI RDMs were calculated using 1-Spearman’s correlation. We then compared the mean fMRI RDM to the mean EEG RDM. The horizontal lines above the curves indicate significant time points (permutation test over stimuli, 10,000 repetitions, FDR-corrected at p < 0.05). The ROIs include occipital-inferior (O-Inf), fusiform (T-LFus), lingual (T-MLing), parahippocampal (T-MTPhip), inferior-temporal (T-IT), and pole-temporal cortex (T-Pole).

We found significant correlations between EEG-fMRI RDMs from about 100 ms onwards, in occipital-inferior, fusiform, and lingual regions, and from about 130 ms onwards in inferior-temporal and pole-temporal regions (for details see Supplementary Table S10). We also observed later representational correspondence between EEG and fMRI starting from about 280 to about 420 ms in the fusiform region, from about 190 ms to about 430 ms in the lingual, and 530 to 560 ms in the inferior temporal region. These results provide an important coordinate to consider how integration methods based on non-invasive EEG and fMRI results perform in comparison to ECoG. For example, we found that occipital inferior, fusiform and lingual were among the first regions to emerge in both ECoG (Figure S2) and EEG-fMRI fusion. According to the ECoG results, parahippocampal cortex was the last region showed selectivity. However, this region showed no significant time point in EEG-fMRI fusion. Moreover, the temporal pole had the longest onset latency in ECoG. This was consistent with EEG-ECoG results which showed that the temporal pole and inferior temporal cortex had longer onset latencies compared to the other regions.

## 5. Discussion

### 5.1. Summary

In this study we investigated the correspondence between EEG and fMRI and ECoG data recorded during object vision in humans at the level of population codes using multivariate analysis methods. Our findings are three-fold. First, the EEG-ECoG comparison revealed a correspondence that drops when viewing-condition invariant representations are assessed. Second, using multivariate pattern analysis, we showed that fMRI-ECoG relation is region dependent; in regions with lower fMRI signal-to-noise-ratio, fMRI-ECoG correlation decreases. Last, comparing the results of EEG-fMRI fusion with that of EEG-ECoG and fMRI-ECoG, we observe both consistencies and inconsistencies across time and region.

### 5.2. Correspondence between EEG and ECoG significantly drops after introducing variations to the stimuli

Our study adds to the large body of research investigation the relationship between EEG and ECoG signals by detailing the relationship in the context of invariant object representations [50, 51]. We observed that when assessing object representations independent of size and orientation, the correlation between EEG and ECoG decoding accuracies was significantly reduced for the chair and the vehicle category. This drop suggests that invariant object representations are located in brain areas that are less accessible by EEG compared to ECoG (Figure 2). This conjecture is further corroborated by the observation that after changing size and orientation of the objects, the EEG classification accuracy decreased, but the ECoG classification accuracy did not (Figure 1).

### 5.3. The impact of signal-to-noise ratio in fMRI on the similarity between fMRI and ECoG

Previous studies on fMRI signal quality comparing different brain regions suggest that fMRI has low signal to noise ratio in temporal regions [26, 44–46]. Consistent with the results, here we observed that in temporal areas of the brain (pole temporal and inferior temporal), the fMRI-ECoG correlation is lower, whereas in regions with higher SNR, such as occipital-inferior and fusiform, the fMRI and ECoG correlation is higher. Together this further highlights the importance of taking the specific sensitivities of different imaging modalities into account when relating and interpreting their results.

### 5.4. Reliability of EEG-fMRI fusion across time and brain regions

When conducting EEG-fMRI representational fusion, i.e. correlating the whole-brain EEG RDMs with the regional fMRI RDMs (i.e. EEG-fMRI fusion [48, 49], we find that consistent with EEG-ECoG RDM correlations across time, occipital-inferior and lingual cortex are among the first regions that emerge around 100 ms. The EEG-fMRI fusion results further suggest that in regions closer to the skull and thus to the EEG electrodes, EEG-fMRI fusion may give more reliable results than in regions further away.

Comparing EEG-fMRI fusion with fMRI-ECoG, we find that pole temporal and temporal inferior fMRI RDMs both had a significant correlation with EEG around 150 ms. However, there was no significant correlation between fMRI and ECoG RDMs for these regions. Pole temporal and temporal inferior are regions typically reported to have lower SNR and more difficult for fMRI to read from [26], which may explain the inconsistency between fMRI-ECoG (Figure 5) and fMRI-EEG (Figure 6) results with regard to these regions.

While ECoG provides high density coverage over a limited cortical region, EEG and fMRI can provide broad coverage over all cortical areas, including deep brain structures in fMRI. This is a key consideration when comparing EEG-fMRI fusion and ECoG results. For areas and time points where EEG and fMRI have reliable readings from the brain and we have enough electrodes from ECoG to read out data, we can expect the best between-modality consistency.

### 5.5. Limitations

One of the study’s limitations is that the EEG-fMRI data were from a different set of participants, compared to that of the ECoG data. Thus, subject-specific signal components could not be assessed. While challenging, future studies may attempt to record EEG, fMRI, and ECoG data from the same participants to be able to do a more direct comparison of the modalities at the level of single individuals.

### 5.6. Conclusion

Our findings of a spatio-temporal correspondence between patterns of category selective responses across ECoG and fMRI/EEG are in line with previous human studies reporting on between-modality correspondence in visual areas [13, 27, 52, 53]. However, these studies did not investigate how the correspondence changes under variations in objects size and/or orientation. Here, we highlighted key differences between EEG-ECoG and fMRI-ECoG, both temporally and spatially, when object variations are introduced. Our study guides interpretation of neuroimaging studies of invariant object recognition when using M/EEG and fMRI by showing when and where we can be more confident about the results.

## 6. Acknowledgment

The EEG and fMRI data were recorded at the Iranian National Brain Mapping Laboratory (NBML). We would also like to thank Morteza Mahdiani and Mahdiyeh Khanbagi for their support and help with some of the data collection and analysis.

