## Supplemental Tables for "A Multivariate Comparison of EEG and fMRI to ECoG Using Visual Object Representations in Humans"

**Table S1. Comparing peak latencies for EEG and ECoG classification time course (for Figure 1A, B, and C).**

| | P-value (EEG peak latency > ECoG peak latency) | mean ± SD (ms) |
|---|---|---|
| **Category selectivity** | 5e-3 | 18 ± 6 |
| **Rotation invariance** | 9.9e-4 | 26 ± 3 |
| **Scale invariance** | 9.9e-4 | 122 ± 41 |

The table shows p-values and mean differences for EEG and ECoG peak latencies when comparing category selectivity, rotation, and scale invariance. P-values were calculated using bootstrap test of 21 participants with 10,000 repetitions.

**Table S2. Comparing correlation coefficients between EEG and ECoG classification time courses across category selectivity, rotation, and scale invariance (for Figure 1D).**

| P-value (category selectivity > rotation invariance) | P-value (category selectivity > scale invariance) | P-value (rotation invariance > scale invariance) |
|---|---|---|
| 1e-4 | 1e-4 | 0.04 |

P-values were calculated using bootstrap test of 21 participants with 10,000 repetitions.

**Table S3. Comparing peak latencies for EEG and ECoG classification time courses (P-value for EEG peak latency > ECoG peak latency, Figure 2A&2B).**

| | **Category** | | | | |
|---|---|---|---|---|---|
| **Classification scheme** | Animal | Chair | Face | Fruit | Vehicle |
| Category selectivity | 1e-4 | 1e-2 | 1e-4 | 9e-1 (n.s.) | 1e-4 |
| Rotation invariance | 5e-1 (n.s.) | 6e-1 (n.s.) | 1e-4 | 5e-1 (n.s.) | 1 (n.s.) |
| Scale invariance | 1e-1 (n.s.) | 6e-2 (n.s.) | 1e-4 | 1e-1 (n.s.) | 9e-1 (n.s.) |

P-values were calculated using bootstrap test of 21 participants with 10,000 repetitions.

**Table S4. Comparing onset latencies for EEG and ECoG classification time courses (P-value for EEG onset latency > ECoG onset latency, Figure 2A&B).**

| | **Category** | | | | |
|---|---|---|---|---|---|
| **Classification scheme** | Animal | Chair | Face | Fruit | Vehicle |
| Category selectivity | 1e-4 | 5e-4 | 7e-4 | 4e-4 | 1e-4 |
| Rotation invariance | 3e-2 | 6e-2 (n.s.) | 2e-3 | 2e-1 (n.s.) | 9e-1 (n.s.) |
| Scale invariance | 1e-2 | 1e-4 | 3e-4 | 5e-4 | 6e-1 (n.s.) |

P-values were calculated using bootstrap test of 16 participants with 10,000 repetitions and signrank test.

**Table S5. P-values for correlation between classification performances of EEG and ECoG for each category (Figure 2B).**

| | Category | | | | |
|---|---|---|---|---|---|
| **Classification scheme** | Animal | Chair | Face | Fruit | Vehicle |
| Category selectivity | 1e-4 | 1e-4 | 1e-4 | 2e-4 | 1e-4 |
| Rotation invariance | 1e-4 | 1e-4 | 1e-4 | 1e-4 | 9e-4 |
| Scale invariance | 1e-4 | 1e-4 | 1e-4 | 1e-4 | 1e-2 |

P-values were calculated using bootstrap test of 21 participants with 10,000 repetitions.

**Table S6. Comparing the peak amplitudes for EEG - ECoG time-resolved point-by-point category-wise correlation (Figure 2C).**

| P-value (category selectivity > rotation invariance) | P-value (category selectivity > scale invariance) | P-value (rotation invariance vs scale invariance_ |
|---|---|---|
| 1e-4 | 1e-4 | 9e-2 |

P-values were calculated using bootstrap test of 21 participants with 10,000 repetitions.

**Table S7. Comparing the onset latencies and peak latencies for EEG - ECoG time-resolved point-by-point category-wise correlation. (Figure 2C).**

| | P-value (category selectivity < rotation invariance) | P-value (category selectivity < scale invariance) | P-value (rotation invariance vs scale invariance) |
|---|---|---|---|
| Onset latency | 2e-3 | 1e-4 | 9e-2 |
| Peak latency | 5e-4 | 4e-4 | 3e-1 |

P-values for peak latencies were calculated using bootstrap test of 21 participants with 10,000 repetitions, and for onset latencies were calculated using bootstrap test of 16 participants with 10,000 repetitions and signrank test.

**Table S8. Peak latencies for whole brain EEG and regional ECOG RDMs comparison (Figure 3B).**

| Region | 95% CI for the first significant peak (ms) |
|---|---|
| Occipital inferior | [104, 121] |
| Lingual | [110, 119] |
| Fusiform | [170, 188] |
| Parahippocampal | [140, 151] |
| Inferior temporal | [171, 199] |
| Pole temporal | No significant peak |

95% confidence intervals were calculated using bootstrap test of 21 participants with 10,000 repetitions.

**Table S9. Onset and peak latencies for regional brain fMRI and ECOG RDMs comparison (Figure 5).**

| Region | Onset latency (ms) | Peak latency (ms) | 95% CI for the first significant peak (ms) |
|---|---|---|---|
| Occipital inferior | 104 | 118 | [112, 132] |
| Lingual | n. s. | No significant peak | No significant peak |
| Fusiform | 76 | 112 | [110, 126] |
| Parahippocampal | n.s. | No significant peak | No significant peak |
| Inferior temporal | n.s. | No significant peak | No significant peak |
| Pole temporal | n.s. | No significant peak | No significant peak |

Onset latencies were calculated using permutation test over stimuli, 10,000 repetitions and FDR-corrected at $p < 0.05$. Peak latencies and 95% confidence intervals were calculated using bootstrap test of 21 participants with 10,000 repetitions.

**Table S10. Onset and peak latencies for regional brain fMRI and EEG RDMs comparison (Figure 6).**

| Region | Onset latency (ms) | Peak latency (ms) | 95% CI for the first significant peak (ms) |
|---|---|---|---|
| Occipital inferior | 90 | 198 | [129:205] |
| Lingual | 90 | 109 | [103:126] |
| Fusiform | 91 | 197 | [164:212] |
| Parahippocampal | n.s. | No significant peak | No significant peak |
| Inferior temporal | 142 | 153 | [142:181] |
| Pole temporal | 135 | 144 | [133:206] |

Onset latencies were calculated using permutation test over stimuli, 10,000 repetitions and FDR-corrected at $p < 0.05$. Peak latencies and 95% confidence intervals were calculated using bootstrap test of 21 participants with 10,000 repetitions.

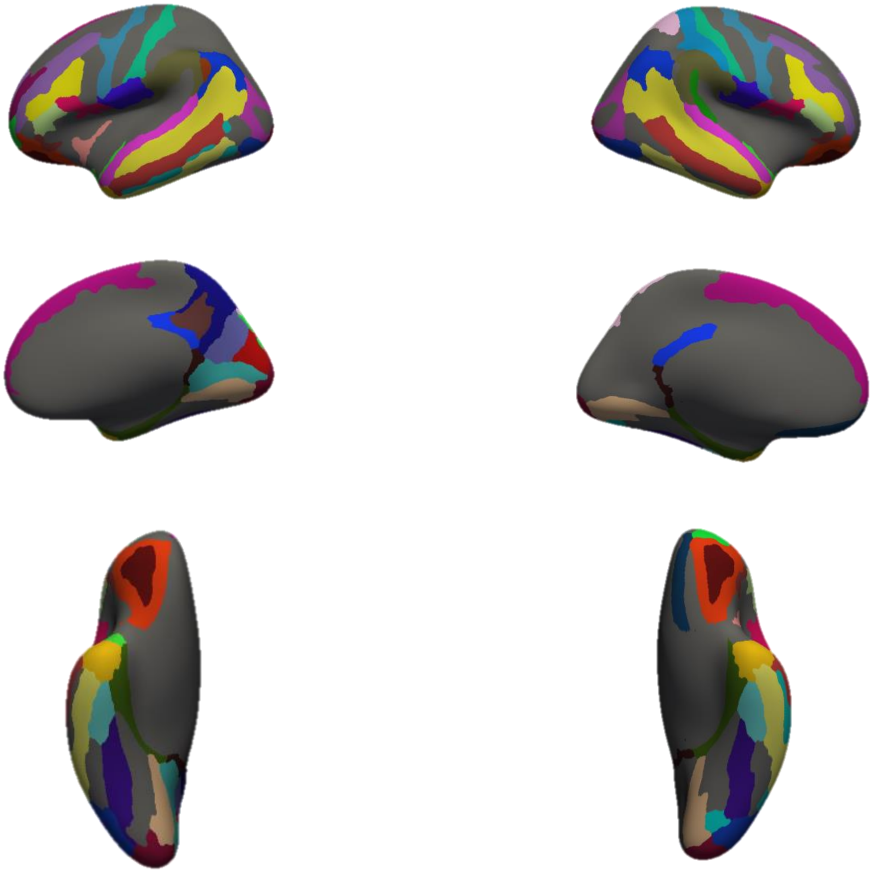

**Figure S1. Colored regions are the regions we used to generate ECoG RDM (for Figure 3A).**

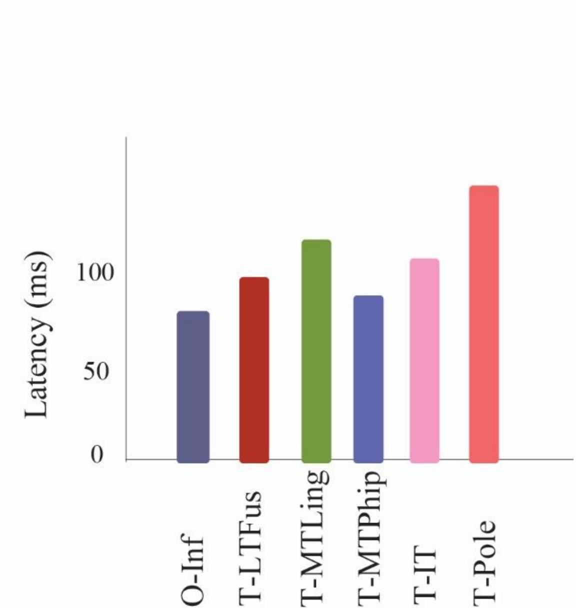

**Figure S2. The mean latencies for ECoG selective electrodes in each location.**
